# A single-strain dropout screen reveals mechanistic links between microbial ecology and metabolism

**DOI:** 10.64898/2026.05.23.727446

**Authors:** Xianfeng Zeng, Xiandong Meng, Allison M. Weakley, Steven Higginbottom, Eugenell Mae Lopez, Ashley V. Cabrera, Ira J. Gray, Brian C. DeFelice, Masahiko Terasaki, Aishan Zhao, Kerrigan R. Hall, Mikhail Levia, Jeanette Arreola, Michael A. Fischbach

**Affiliations:** Department of Bioengineering, Stanford University, Stanford, CA 94305, USA; ChEM-H Institute, Stanford University, Stanford, CA 94305, USA; Chan Zuckerberg Biohub, San Francisco, CA 94158, USA

## Abstract

The complexity of the gut microbiome has made it challenging to define the role of individual species in community-level function. Here, we constructed 56 single-strain dropout variants of a defined 118-member community and used each one to colonize a group of germ-free mice. In many cases, removing a single strain triggered a large reordering of a small group of species, which in turn altered the community’s metabolic output. *En bloc* removal of the eight-strain acetogen compartment markedly reduced acetate production and caused intestinal H_2_ accumulation and bloating; a specific subset of four acetogens was sufficient to relieve bloating and restore acetate production. Together, these data show that small disturbances in community composition can trigger a confined ecological reorganization with a large chemical phenotype, and they reveal novel strategies for engineering communities with altered metabolic output.

## INTRODUCTION

The gut microbiome impacts a variety of host phenotypes including the response to checkpoint inhibitors^1–6^, the efficacy of vaccination^7^, nutrient extraction^8^, and colonization resistance to enteric pathogens^9–11^. However, in humans and conventionally colonized mice, it can be difficult to identify which strains are involved in a phenotype of interest and by what mechanisms they act. In biological systems, the most common way to figure out which components are involved is to remove one at a time (i.e., a test of necessity). In model organisms like mice, worms, or yeast, single-gene knockouts have long served as a foundational tool for uncovering mechanisms. But until recently, the same logic was not applicable to the microbiome. Model communities were either complex but undefined, making it impossible to isolate the role of any one strain; or defined but overly simplified, limiting physiological relevance.

To address this gap, we recently developed a model system for the gut microbiome, hCom2, that is complex and defined. In principle, a system like this could be used to identify strains involved in host phenotypes of interest. However, we do not yet know—in general terms—how such a system will respond to the removal of one or several strains^12^. Here, as a starting point for this kind of work, we constructed 56 single-strain dropout variants of hCom2, focusing on the Firmicutes, which are important metabolically and immunologically but not as well characterized as the Bacteroidales. We colonized germ-free mice with each of these communities and characterized them in two ways: high-resolution metagenomic sequencing, to assess how each dropout affects ecological structure; and targeted metabolomic analysis, to identify the chemical consequences of eliminating individual strains.

We observed variable ecological consequences across the dropout communities. In some cases, strain removal had little or no effect on community architecture; in others, it triggered a substantial reorganization. Strain dropout communities frequently exhibit metabolic phenotypes, often because of a dropout-triggered reorganization (i.e., strain removal causes another strain to change in abundance, which in turn leads to a shift in metabolite production). Thus, strain removal is an effective way to reveal phenotypes because it unleashes ecological events that have a large effect size but are limited to a small clique within the community and therefore lend themselves to mechanistic inquiry.

We characterized one metabolic group in detail: the acetogens. By colonizing germ-free mice with a community lacking all acetogenic strains, we observed that the absence of acetogens led to a marked reduction in acetate production and intestinal bloating due to a buildup of hydrogen gas. Some subsets of acetogens rescue acetate production and mitigate gas buildup while others had little effect, indicating that strain contributions to this niche are unequal.

### Design of the single-strain dropout screen

In previous work, we developed a 119-strain community that serves as a model system for the human gut microbiome, hCom2^13^. In this study, we used a closely related variant (hCom2a, 118 strains) in which *Ruminococcus albus* and *Ruminococcus flavefaciens* were omitted and *Lactobacillus plantarum* was added to facilitate studies of bile acid metabolism [*Zeng et al,* 2026b].

Our goal was to perform a screen in which we would omit one strain at a time from hCom2a as a starting point for defining the ecological and biochemical role of each strain in the community. The Bacteroidetes are well known for their role in harvesting dietary and host-derived polysaccharides^14–20^, so we focused instead on the Firmicutes. Although they are known to play a role in immune modulation^21–24^, metabolite production^25–28^, and colonization resistance^9,10,29^, less is known about strain-level contributions and mechanisms.

Our goal was to generate 68 single-strain dropouts: all 63 Firmicutes in hCom2a and several strains outside of the Firmicutes that were of metabolic interest: *Bilophila wadsworthia* ATCC 49260, *Bacteroides dorei* DSM 17855, *Intestinimonas butyriciproducens* DSM 26588*, Eggerthella lenta* DSM 2243, and *Akkermansia muciniphila* ATCC BAA-835.

### Validating the single-strain dropout communities

We set out to construct 68 single-strain dropout variants of hCom2. We started by reviving frozen stocks of the constituent strains in liquid culture. To construct each dropout community, we pooled all but one of the strains, omitting the strain of interest. Communities were assembled and assayed in batches of 6-12. We colonized 3-5 germ-free Swiss-Webster mice with each community by oral gavage with 200 µL of mixed culture, administered twice on successive days to ensure stable colonization. Two cages of hCom2a-colonized mice were typically included in each batch as a control (**fig. s1a-c**). Male and female mice were both used; each dropout community was tested in one or the other. We observed that community architecture was consistent in male and female mice gavaged with the same inoculum (**fig. s1d-e**).

Four weeks after colonization, we sacrificed the mice and collected cecal contents, small intestinal contents, and urine. To test whether the strain we intended to omit was indeed absent, we profiled community composition in the inoculum and cecal contents by high-resolution metagenomic analysis, with read-level mapping performed using NinjaMap. We tested the validity of each dropout by confirming the absence of the targeted strain in cecal metagenomic data. Of 68 attempted dropouts, 56 succeeded and 12 failed; all downstream analyses focused on these 56 communities (**Fig. 1b**, **fig. s1f**).

**Figure 1:**
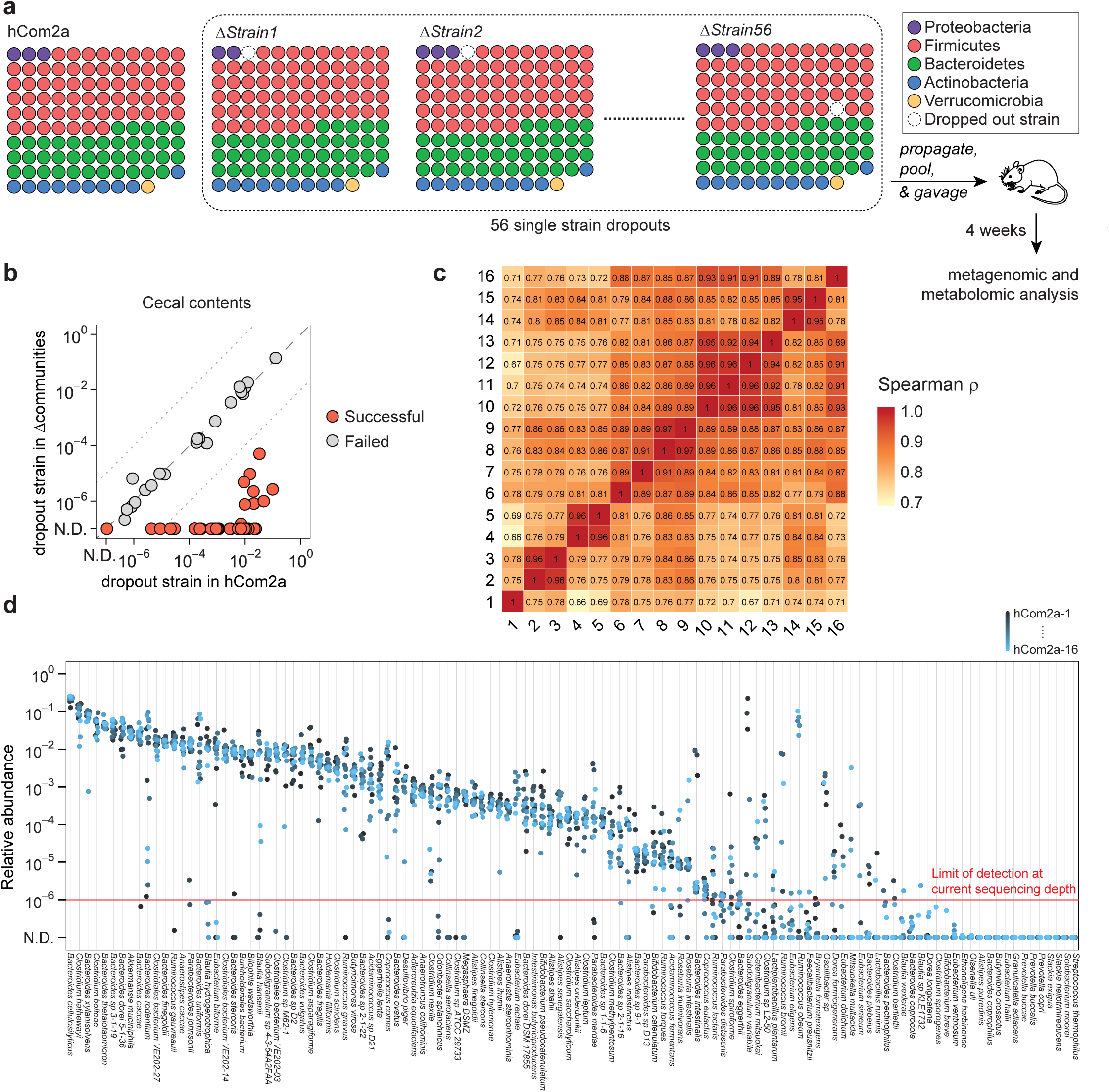
A single strain dropout screening within a defined community. (a) Schematic of the dropout screen. Frozen stocks of each strain in the community minus the strain to be dropped out were thawed, propagated in growth medium, mixed in equal volume, and used to colonize germ-free C57BL/6N mice (n = 3–6 mice per replicate) by oral gavage. After 4 weeks of colonization, mice fed ad libitum on a chow diet were sacrificed. Intestinal contents were collected, subjected to metagenomic sequencing, and analyzed by NinjaMap to measure the composition of the community. Cecal contents and urine were collected and analyzed by LC-MS to profile a panel of microbiome-derived metabolites. Dropouts were performed in 16 experimental batches. Each batch compared the parental hCom2a community to a subset of single-strain (b) We constructed 68 single-strain dropout communities, of which 56 were successful. The plot compares the relative abundances of each strain in the dropout community (y-axis) to the corresponding hCom2a control community (x-axis). We defined a >100-fold reduction in the omitted strain as a successful dropout; dropout communities failing this criterion were excluded from subsequent analysis. (c) Inter-batch consistency of control communities. Heatmap of pairwise Spearman correlation coefficients based on strain-level relative abundances across all hCom2a control mice from different experimental batches. (d) Composition and strain-level relative abundances of all hCom2a control groups. Each column represents the abundance of a single hCom2a strain across 16 control mice from different batches. The red vertical line indicates the approximate limit of detection (∼10^-6^) based on a sequencing depth of 20–30M reads/sample.

hCom2a composition across batches (i.e., biological replicates) was reproducible: 52/102 strains (51%) were tightly distributed (SD < 0.5 log_10_ units, ∼3-fold variation), and 71/102 (70%) were broadly consistent (SD < 1 log_10_ units, ∼10-fold variation). The remaining 31 strains had some degree of variability; we speculate that this may have resulted from inconsistent success in strain revival or biological fluctuations in relative abundance (**Fig. 1c-d**). 16 strains were below the limit of detection in all hCom2a groups. This does not necessarily imply that they are absent, as low-abundance strains may evade detection due to sampling depth or stochastic variation.

### Compartmentalized reordering in response to single-strain dropout

Next, we sought to understand how the community responds ecologically to the removal of an individual strain. Given the presumed degree of redundancy, the removal of one strain could go unnoticed by the rest of the community. Alternatively, in light of the rich web of presumed ecological interactions, omitting an important strain could cause a major restructuring of the ecosystem^30^.

To determine the ecological consequences of strain removal, we analyzed high-resolution metagenomic sequencing data to determine which strains change significantly in each dropout community, using an approach that accounts for each strain’s distribution of relative abundances across samples (see **Methods**, **fig. s2a**). In most single-strain dropout communities, a precise ecological perturbation was observed: a large majority of the strains were unaffected, while a small subset—typically one to four—shifted markedly in relative abundance (**Fig. 2**, **fig. s2b-c**). A smaller number of dropouts had a more extreme profile: for a subset of strains (*A. fermentans*, *D. longicatena*, *C. bartlettii*, *C. comes*), omission had no significant impact on community composition, whereas removing *R. gauvreauii* or *C. asparagiforme* led to profound ecological restructuring. In total, 65% of the strains in hCom2a (77/118) respond to at least one dropout. Together, these results reveal that removing a single strain often unleashes an ecological reordering that is large in magnitude but limited to a small set of species within the community.

**Figure 2:**
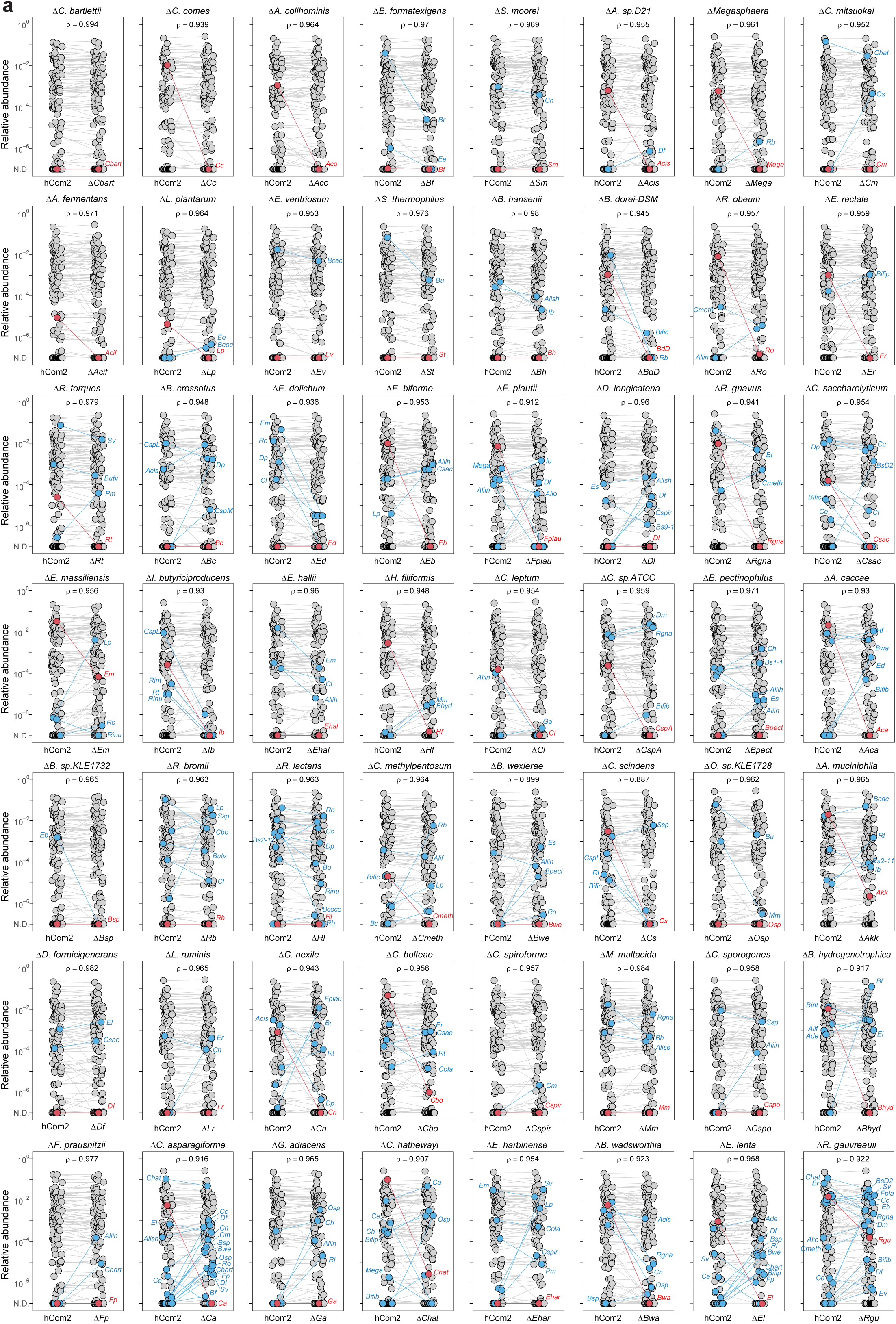
The ecological impact of single strain removal varies across communities. (a) Median relative abundances of hCom2a-colonized mice (hCom2a) compared to single-strain dropout communities (ΔStrain). Each panel represents one dropout community; each dot is an individual strain, and the collection of dots in a column represents the community averaged over 3-6 mice from the same community. The omitted strain is highlighted in red; blue dots highlight strains whose relative abundance differs significantly between hCom2a and the dropout community (|Z| > 2). Panels are ordered by ecological impact, defined as the variance of Z-scores resulting from the removal of a single strain. For strain name abbreviations, see Table s1e.

### Strain dropouts influence community-level metabolism

To determine whether dropping out individual strains has an impact on community-level metabolism, we performed a targeted metabolomic analysis of cecal contents and urine from community-colonized mice at the time of sacrifice. We focused on metabolites that are produced exclusively or predominantly by the microbiota, measuring short-chain fatty acids (SCFAs) and bile acids in cecal contents and quantifying 39 additional microbiome-derived metabolites in urine (**Table S3**). We then analyzed, for each metabolite, which dropout communities led to a significant increase or decrease in production.

In spite of the fact that most microbiome-derived metabolites are produced by more than one organism in hCom2a^12^ (**Fig. 3a**), several of the single-strain dropout communities exhibit a marked shift in metabolite production. In five such examples covering nine strain dropouts, the accompanying sequencing data offers a hypothesis about the ecological mechanism underlying the metabolic phenotype.

**Figure 3:**
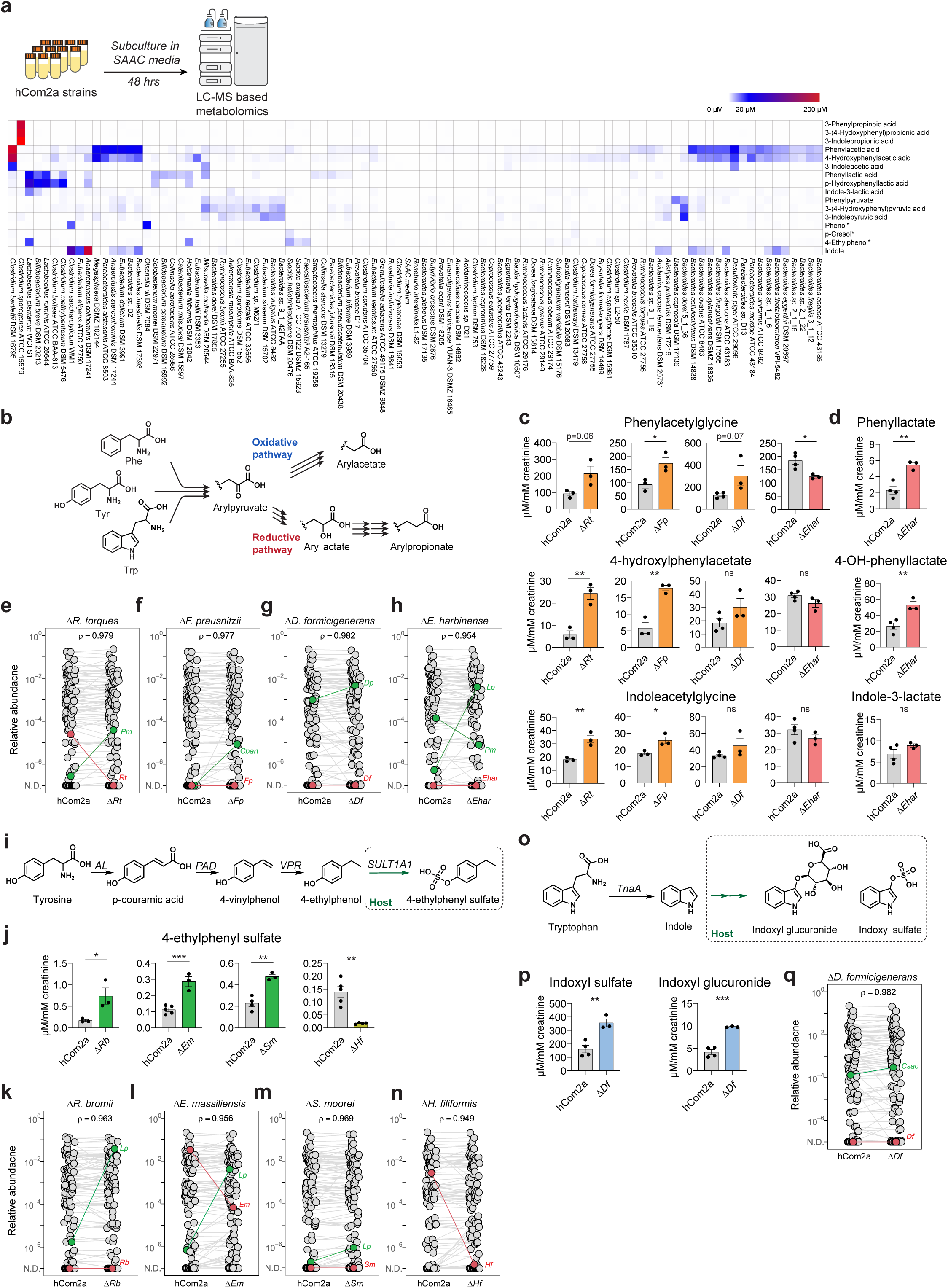
Strain removal impacts community-level metabolism. (**a**) *In vitro* profiling of aromatic amino acid (AAA) fermentation. (Top) Experimental schematic. (Bottom) Heatmap of AAA fermentation products from selected hCom2a strains. Phenol, 4-cresol, and 4-ethylphenol production were assessed following the addition of specific precursors (4-hydroxybenzoic acid, 4-hydroxyphenylpyruvic acid, and 4-vinylphenol, respectively); other metabolites were profiled in base SAAC medium. (**b**) Schematic of oxidative and reductive AAA fermentation pathways. (**c-d**) Impact of strain removal on oxidative AAA fermentation. (**c**) Urinary levels of oxidized AAA fermentation products. Mice colonized by three dropouts (*ΔRt, ΔFp, ΔDf*) showed increased levels of oxidized AAA metabolites, while *ΔEhar*-colonized mice showed decreased levels. (**d**) Urinary AAA-lactic acid levels in hCom2a versus *ΔEhar*. (**e-h**) Changes in community composition in *ΔRt, ΔFp, ΔDf*, and *ΔEhar*. (**e**) Median relative abundances of hCom2a vs *ΔRt* in cecal contents. Each dot represents an individual strain; dots in a single column correspond to the median abundances from 3–4 co-housed mice (one cage). The omitted strain (*Rt*) is highlighted in red. Green dots indicate potential contributors to the altered metabolic phenotype. Similar data are shown for *ΔFp* (**f**), *ΔDf* (**g**), and *ΔEhar* (**h**). (**i-n**) 4-ethylphenol production. (**i**) Schematic of 4-ethylphenol biosynthesis. AL: ammonia lyase; PAD: phenolic acid decarboxylase; VPR: vinyl phenol reductase. (**j**) Urinary 4-ethylphenol levels across dropout communities. (**k-n**) Relative abundance analysis (as in (**e**)) for *ΔRb, ΔEm, ΔSm, and ΔHf*. (**o-q**) Indole production. (**o**) Schematic of indole biosynthesis from tryptophan via tryptophanase (TNA), and the subsequent conversion of indole to indoxyl sulfate and indoxyl glucuronide by the host. (**p**) Urinary indole derivatives in hCom2a-versus *ΔDf-*colonized mice. (**q**) Relative abundance analysis (as in (**e**)) for *ΔDf*. All graphs show mean +/-SEM. Statistical significance was determined using a two-tailed Student’s t-test; p < 0.05 (*), p < 0.01 (**), p < 0.001 (***).

### Oxidative aromatic amino acid metabolism

We observed a substantial increase in the production of oxidative AAA metabolites in mice colonized by the *Ruminococcus torques* (Δ*Rt*)*, Faecalibacterium prausnitzii* (Δ*Fp*), and *Dorea formicigenerans* (Δ*Df*) dropouts (**Fig. 3b-c, fig. s3c**). To gain insight into the ecological underpinnings of this phenotype, we compared the list of strains that change significantly in each dropout (**Fig. 3e-g**) with results from an experiment in which we profiled AAA metabolite production by each strain individually *in vitro* (**Fig. 3a**). In each dropout, we found that one of the few strains that increased in relative abundance was a producer of oxidized AAA metabolites (*P. merdae* in Δ*Rt,* Z = 2.19; *C. bartlettii* in Δ*Fp, Z > 5; D. piger in* Δ*Df,* Z = 1.04), suggesting the possibility that the effect on the AAA metabolite pool was mediated by an ecological interaction between the omitted strain and a (distinct) metabolizing strain.

In a fourth dropout community, we observed a reduction in oxidative AAA metabolism, accompanied by an accumulation of aryllactate—an intermediate of reductive AAA metabolism (**Fig. 3c-d**). High-resolution metagenomic analysis of the cecal contents revealed a decrease (Z = -2.58) in the oxidative AAA-producing strain *P. merdae* (which had bloomed in Δ*Rt*), and an increase in *L. plantarum* (Z > 5). Notably, in other communities where *L. plantarum* bloomed, aryllactate production did not increase, suggesting that the shift in metabolite production was not driven by *L. plantarum* alone. Instead, the combination of *P. merdae* depletion and *L. plantarum* enrichment appears to redirect aromatic amino acid metabolism from oxidative to reductive AAA metabolism.

### 4-ethylphenol metabolism

4-Ethylphenyl sulfate was elevated in mice colonized by three dropout communities—Δ*Rb* (*Ruminococcus bromii*), Δ*Em* (*Eisenbergiella massiliensis*), and Δ*Sm* (*Solobacterium moorei*)—but was absent in the Δ*Hf* (*Holdemania filiformis*) dropout (**Fig. 3i-j, fig. s3d**). To investigate potential contributors, we examined the sequencing data and cross-referenced strains known to produce 4-ethylphenol *in vitro*. In all three communities in which 4-ethylphenyl sulfate increased, we observed an enrichment of *L. plantarum*, one of the most prolific 4-ethylphenol producers *in vitro* (**Fig. 3k-m**). In contrast, the diminution of 4-ethylphenyl sulfate in Δ*Hf* likely reflects the loss of *H. filiformis* itself, a known 4-ethylphenol producer (**Fig. 3n**).

### Indole production

A similar observation was made in Δ*Df*-colonized mice: the level of the indole-derived metabolites indoxyl sulfate and indoxyl glucuronide increased (**Fig. 3o-p**). In the dropout mice, we observed a higher relative abundance of *C. saccharolyticum* (**Fig. 3q**); notably, the increase was modest (∼2-fold) but highly significant (Z = 2.20), as *Csac* typically colonizes in a very tight relative abundance range. *Csac* is one of only two strains in the community that convert tryptophan to indole robustly (**Fig. 3a**), consistent with a model in which an ecological interaction leads to an increase in metabolite production.

Taken together, these results demonstrate that single-strain perturbations induce discrete, mechanistically interpretable shifts in community-level metabolism. Notably, we identify three modes by which a perturbation impacts the chemical output of a community. (*i*) Most commonly, removal of strain 1 leads to the expansion of strain 2, a metabolite producer (e.g., *L. plantarum* or *P. merdae*), redirecting flux through strain 2’s pathway. (*ii*) Less commonly, removal of a metabolite-producing strain (e.g., *H. filiformis*) leads directly to a reduction in the corresponding metabolite. (*iii*) In rare cases, a stochastic change in colonization enables a low-abundance, metabolite-producing strain to bloom, leading to a durable shift in community metabolism, even though the strain roster is identical at our level of resolution (**fig. s3f-m, supplementary text**). These findings underscore that the gut microbiome does not function as a monolithic, redundant system, but rather as a collection of metabolic modules that can be rewired by ecological rearrangements.

### Systematic dissection of the acetogen compartment in hCom2a

The examples above show that the chemical output of the microbiome can be altered by strain-level perturbations. However, acetate—the most abundant metabolic product of the gut microbiome—stood out as an exception: none of our dropout communities had significantly altered acetate levels (**fig. s3n**).

This is perhaps unsurprising: in a case of extreme metabolic redundancy, 113 of 117 strains (96.6%) in hCom2a encode a complete EMP pathway together with the canonical Pta-AckA cassette for converting pyruvate to acetate (**Table S4**). However, acetate can also be produced by the Wood-Ljungdahl pathway (WLP, **Fig. 4a**), in which acetogenic Firmicutes convert CO₂ and H₂—major byproducts of fermentation—directly into acetate. This raises a question that goes beyond species-level redundancy: do these distinct acetate-producing compartments compensate for each other, such that total acetate output is fixed? Or is the community-level output of acetate determined by each pathway independently? To address this question, we focused on acetogens as a tractable entry point for perturbing acetate production while leaving the EMP compartment largely intact.

**Figure 4:**
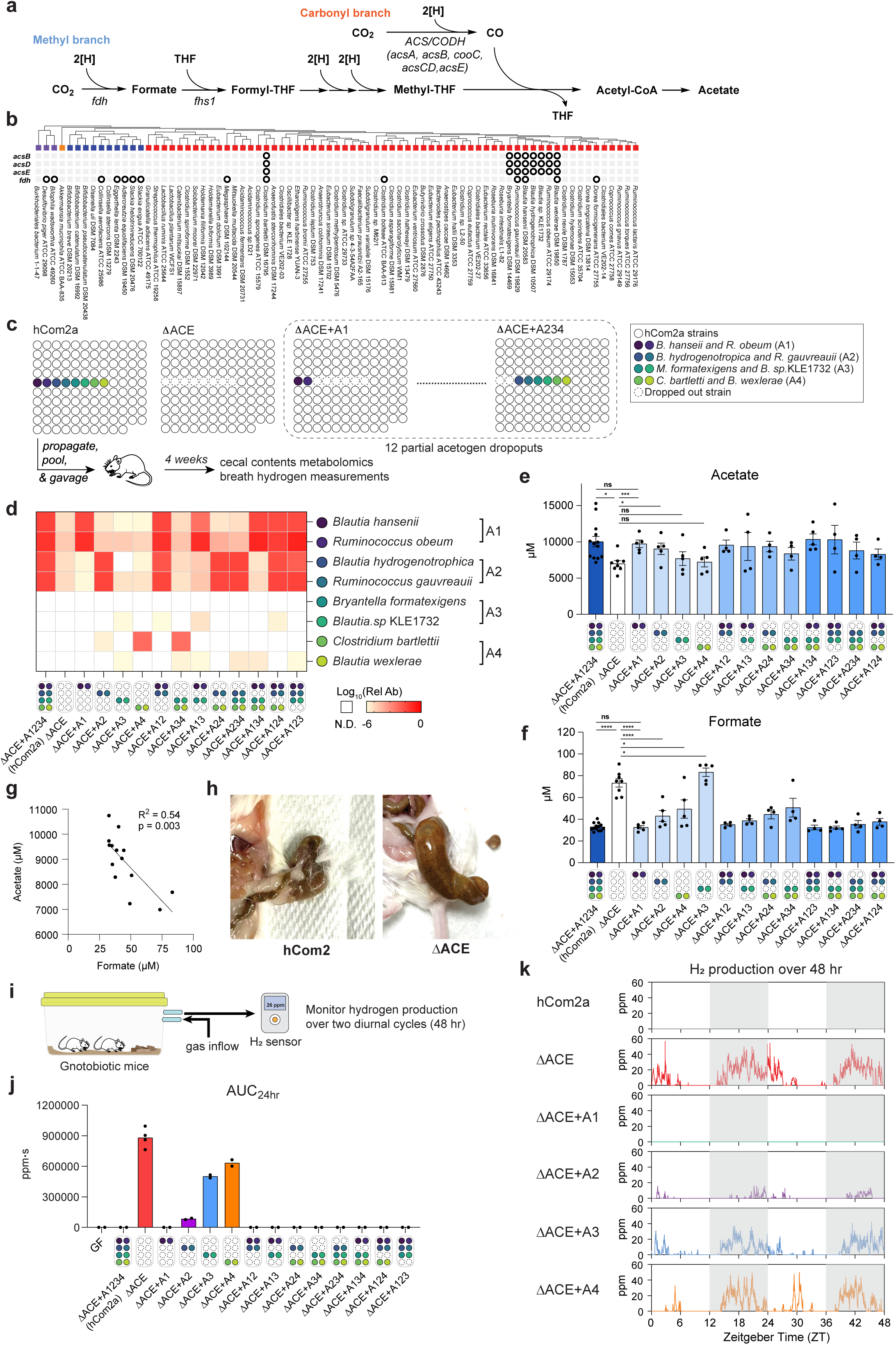
Systematic dissection of the acetogen niche in a defined community. (**a**) Schematic representation of the Wood-Ljungdahl pathway (WLP). (**b**) Distribution of genes encoding core WLP enzymes and formate dehydrogenase in bacterial species from hCom2a. Tetrahydrofolate-related enzymes (e.g., *fhs*) are widely present and were excluded from the analysis. Black circles denote the presence of a gene in the family indicated by each row. No hits were detected in Bacteroidetes, which are omitted in the figure. (**c**) Experimental strategy for dissection of the acetogen compartment. Acetogens in hCom2a were classified into four groups (A1–A4), A1: *B. hansenii*, *R. obeum*; A2: *R. gauvreauii*, *B. hydrogenotropica*; A3: *M. formatexigens*, *B. sp. KLE1732*; A4: *C. bartlettii*, *B. wexlerae.* A complete acetogen-dropout community (*Δ*ACE) was generated by removing all eight acetogens. Partial acetogen-dropout communities were generated by adding back specific acetogen groups to *Δ*ACE (e.g., *Δ*ACE *+* A12 indicates that groups A1 and A2 were restored). All communities were used to colonize germ-mice by oral gavage. After 4 weeks of colonization, mice were sacrificed and analyzed by sequencing and mass spectrometry. Colored dots denote acetogen strains included in each group (see legend for strain identity). (**d**) Cecal colonization of acetogens. The heatmap shows the relative abundances of acetogens in hCom2a and the dropout communities; values below the limit of detection (<10^-6^) are colored white. (**e-f**) Targeted metabolic profiling of acetate (**e**) and formate (**f**) in cecal contents. (**g**) The levels of acetate and format in cecal contents are negatively correlated across community groups. (**h**) Intestinal gas accumulation in *Δ*ACE mice. Representative image of cecal contents from hCom2a- and *Δ*ACE-colonized mice, illustrating marked gas accumulation in *Δ*ACE-colonized mice. (**i-k**) Continuous hydrogen monitoring. (**i**) Schematic of the gnotobiotic gas-monitoring system. Exhaust air from sealed cages was routed to a passive-diffusion sensor for diurnal monitoring. (**j**) Area under the curve (AUC) of H_2_ production over 24 h. (**k**) Diurnal H_2_ production profiles. Continuous 48 h measurements from cages colonized with the indicated communities. Shaded areas indicate the dark (active) cycle. Zeitgeber Time (ZT) 0 corresponds to lights on. Note the correlation between peaks and typical feeding behavior. All graphs show mean +/-SEM. Statistical significance was determined using two-tailed Student’s t-test; p < 0.05 (*), p < 0.01 (**), p < 0.001 (***).

Acetogens can be identified computationally by the presence of genes encoding the WLP, which consists of two branches: a methyl branch, in which CO_2_ is reduced to a methyl group via the formate–methyl-THF route; and a carbonyl branch, in which CO_2_ is reduced to CO by carbon monoxide dehydrogenase (CODH), then combined with the methyl group by acetyl-CoA synthase (ACS) to form acetyl-CoA. We started by using BLAST to determine which strains in hCom2a harbor the WLP, searching their genomes for genes encoding the ACS/CODH complex. We identified eight strains that harbor the corresponding gene cluster (**Fig. 4b**) and focused our experimental efforts on these strains.

Next, we constructed a community lacking all eight acetogens (ΔACE). In parallel, we constructed a series of partial acetogen dropouts by grouping them into four pairs and reintroducing them in twelve sets of two, four, or six strains. Mice were colonized with these communities for four weeks, followed by metagenomic and metabolomic profiling of cecal contents. Sequencing data confirmed that the strains we intended to drop out were indeed omitted (**Fig. 4c-d**).

The complete removal of all acetogens (ΔACE) led to a marked shift in cecal metabolites: acetate levels dropped from ∼10 mM to ∼7 mM, while the formate concentration doubled (**Fig. 4e-f, fig. s4a-bd**). Although this has never been demonstrated before, it is consistent with a prior estimate, from *ex vivo* isotope tracing, that the WLP accounts for one-quarter to one-third of the total fecal acetate pool^31^. Propionate and butyrate were unaffected, consistent with a perturbation to the SCFA pool due entirely to the loss of the WLP pathway. Acetate production was restored by two of the strain pairs that colonized robustly, while the other two—which colonized less efficiently—did not complement the dropout phenotype. Together, these results suggest that as few as two species are sufficient to carry out full-scale acetogenesis, provided they colonize efficiently. Of note, *C. bartlettii*—typically a low-abundance colonist—reached higher abundance in the absence of more robust acetogens, suggesting competitive interactions within the niche.

In the absence of CODH/ACS-expressing bacteria, CO_2_ can be reduced to formate by non-acetogens that harbor formate dehydrogenase. Across all communities, acetate and formate levels were strongly anti-correlated (**Fig. 4g**), indicating that acetogen deficiency diverts one-carbon flux from the WLP into formate.

### Acetogenesis regulates intestinal gas homeostasis

In acetogen-deficient mice (ΔACE), we observed a pronounced accumulation of intestinal gas, a phenotype absent in mice colonized with the parental community (hCom2a) (**Fig. 4h**). The WLP pathway consumes hydrogen (H₂) and carbon dioxide (CO₂) to generate acetate, acting as a gas sink. We reasoned that the accumulation of gas was caused by the loss of WLP pathway activity.

To test this hypothesis, we measured expelled hydrogen gas using a custom “mouse hotel” system (**Fig. 4i, fig. s4c**). In this setup, two gnotobiotic mice were co-housed in a sealed cage; the cage exhaust was continuously routed through a hydrogen gas sensor. Because hydrogen production can fluctuate with feeding, gas production was recorded continuously across a diurnal cycle. While individual biological replicates showed distinct patterns, the total hydrogen output was consistent (**fig. s4d-e**).

As expected, hCom2a-colonized mice generated no detectable hydrogen gas, consistent with complete consumption of fermentative H₂ by the WLP (**Fig. 4j-k**). In contrast, ΔACE-colonized mice produced a large quantity of H_2_, predominantly during the dark (feeding) phase. These data suggest that the gas buildup in acetogen-deficient mice is composed in part of hydrogen that would ordinarily be consumed by the WLP.

Add-back experiments revealed functional inequality among acetogens. The two acetogen pairs that colonize efficiently largely rescued the hydrogen accumulation phenotype, while the low-efficiency acetogen colonists failed to do the same. Of note, combining the two poorly colonizing groups (ΔACE + A34) restored H_2_ to an undetectable level, suggesting that multiple weak acetogens can function additively to consume the available hydrogen gas. In all communities containing four or more acetogens (two or three groups added back), hydrogen levels were undetectable, indicating that a minimal threshold of acetogen colonization is sufficient to restore WLP-dependent gas balance (**Fig. 4j-k, fig.s4f**). Communities with detectable levels of hydrogen production (ΔACE, ΔACE + A2, ΔACE + A3, ΔACE + A4) had higher cecal formate and lower acetate levels, demonstrating that H_2_ consumption and acetate production are coupled through the WLP.

## DISCUSSION

The ability to remove individual strains one at a time—previously not feasible to do at scale—has revealed rich insights into community ecology. At the outset, we had one major concern about the design of this study: the microbiome is so complex that dropping out a single strain would rarely, if ever, yield a measurable phenotype; other strains would compensate, masking the phenotype^12^.

Our concerns about phenotypic masking were misplaced, in large part because the consequences of strain removal were unexpected. While some dropouts had a loss-of-function phenotype related to the strain in question, more often the dropout triggered an ecological reordering. In these cases, the loss of one strain led to the bloom of another, which in turn altered the community’s metabolic output. If this pattern proves generally true, it would imply that seemingly minor strain-level differences among individuals could have an outsize impact on the chemical output of their gut communities. It would also have important implications for engineering; the best way to boost the level of a metabolite producer might be to eliminate a strain impeding its colonization.

We were unable to repeat each condition in the single-strain dropout experiment. As a result, we view the results as a screen; they are consistent with the possibility of a phenotype, but each one will need to be repeated before concluding that the ecological sequelae are real (i.e., the absence of strain 1 leads to a change in the relative abundance of strain 2). We expect that some of these phenotypes will be reproducible and others will not. However, we are more confident about the relationships we observed between strain 2 and the resulting metabolic phenotypes.

Intestinal bloating and gas accumulation are often attributed to the overgrowth of hydrogen-producing microbes^32,33^. Our data suggests a different possibility: some instances of bloating might result from the absence of acetogens. In hCom2, the absence of acetogens led to a profound accumulation of hydrogen gas, resulting in a visible bloating phenotype. In other communities, two additional classes of hydrogenotrophs—sulfate-reducing bacteria and methanogens—might play an important role as well.

Together, these results illustrate the power of using defined communities and combinatorial dropout/addback strategies to dissect functional contributions within complex microbial communities. We were able to map, with strain-level resolution, the role of acetogens in consuming intestinal H_2_ and synthesizing acetate. While all eight acetogens harbor the Wood–Ljungdahl pathway, their contributions to hydrogen assimilation are unequal; a minimal set of four acetogen species are required to robustly maintain gas balance in the gut. These results shed light on the mechanistic underpinnings of intestinal bloating, and they suggest a new strategy for treating IBS based on the restoration of acetogens.

## ACKNOWLEDGMENTS

We are deeply indebted to members of the Fischbach laboratory for helpful discussions and suggestions, especially Min Wang and Alice Cheng for microbiology input, Thao Pham for help with culturomics, and Maria Prado, Amanda Espinoza and Joanne Au for keeping the laboratory running. X.Z. is a Fraternal Order of Eagles Fellow of the Damon Runyon Cancer Research Foundation, DRG-2517-24. This work was supported by the Helmsley Foundation, the DOW (Vannevar Bush Faculty Fellowship), NIH (DK101674), NSF (2125383), and the Stanford Microbiome Therapies Initiative.

## AUTHOR CONTRIBUTIONS

X.Z. and M.A.F. conceived the study, X.Z. performed most experiments, including data analysis, community construction, cloning, and method development. X.M. helped with bioinformatics, A.M.W, A.V.C. performed metagenomic sequencing, S.H., E.M.L., assisted with gnotobiotic experiments, I.J.G., B.C.D. helped with metabolomics, M.T., A.Z., K.R.H., M.L, and J.A. helped with community preparation. The manuscript was drafted by X.Z. and M.A.F. and reviewed and edited by X.Z. and M.A.F.

## SUPPLEMENTARY MATERIALS

Materials and Methods

Figures s1-s4

Table s1-s4

## MATERIALS AND METHODS

### Bacterial strains and culture conditions

All bacterial strains used in the synthetic community were sourced from the American Type Culture Collection (ATCC), the Leibniz Institute DSMZ–German Collection of Microorganisms and Cell Cultures GmbH (DSMZ), BEI Resources (BEI), and additional sources as specified. Strains were cultured in one of the following media: modified Columbia medium, modified Peptone Yeast Extract Glucose (PYG) medium, Yeast Casitone Fatty Acids (YCFA) medium, modified Gifu medium, modified Baar medium, bifidobacterium medium, ethanoligenens medium, or chopped meat medium. Where necessary, specific nutrients were added to individual cultures to support optimal growth (see **Table S1**).

### Preparation of synthetic community

For all community culturing, strains were cultured in anaerobic conditions (10% CO_2_, 5% H_2_, and 85% N_2_). For strain purification and storage, each strain was streaked on its respective agar (see **Table S1**), assessed for purity, and a single colony was propagated in 200-300 mL of the corresponding liquid medium. Cultures were harvested at early stationary phase, centrifuged, and resuspended in 25% glycerol prepared in the same growth medium. Aliquots of 300 μL were stored in individual matrix tubes at –80 °C.

To prepare synthetic communities for *in vivo* experiments, frozen stocks of each strain were thawed and propagated in 40 mL of their respective growth medium in 50 mL Falcon tubes for 72 h. Equal volumes of each culture comprising the intended community (including hCom2a and strain dropout communities) were pooled, centrifuged at 4,700 × g for 30 minutes, washed, and resuspended in an equal volume of 25% glycerol in modified Columbia medium. The final synthetic community mixture was aliquoted into 700 μL portions and stored at –80 °C until use.

### Gnotobiotic mouse experiments

Germ-free Swiss-Webster or C57BL/6N male mice (6–8 weeks of age) were originally obtained from Taconic Biosciences (Hudson, NY). Colonies were maintained in gnotobiotic isolators and provided autoclaved chow and water *ad libitum*. All animal procedures were approved by the Stanford University Institutional Animal Care and Use Committee (IACUC).

For colonization, 1.2 mL glycerol stocks of synthetic communities were thawed and mixed thoroughly at room temperature. Mice were orally gavaged with 200 μL of the mixed culture. To ensure robust colonization by all strains, mice were gavaged once daily for two consecutive days in all experiments.

The mice were maintained on a standard diet (LabDiet 5k67; 0.2% Trp) for 4 weeks (unless otherwise stated) before sacrifice (fed *ad libitum*). Mice were euthanized humanely by CO_2_ asphyxiation and, urine, liver, plasma the luminal contents of the small intestine, cecum, and colon were collected at the same time of day and stored at –80 °C until use.

### Measurement of hydrogen during a diurnal cycle

To monitor breath hydrogen production by mice over a diurnal cycle, individual mouse cages (Isocage positive) were modified to allow redirection of cage exhaust air through a sealed tubing system. The adapter was secured through parafilm and a 3M tape. The tubing was connected to a hydrogen sensor (Forensics detectors, FD-90A-H2) capable of detecting real-time changes in hydrogen concentration. The entire setup, including the modified cage, tubing, and sensor—was assembled and placed in the animal facility overnight under standard housing conditions (**fig. s4c**). Sensor readings (ppm) were recorded continuously using an action camera (DJI Action 4) aimed at the display; a red headlamp (Black diamond Storm-500R) provided low-disturbance illumination during the dark phase. For validation, three independent cages colonized with *Δ*ACE were monitored for 24 h; area under the curve (AUC, ppm·s) over 24 h was used as the total hydrogen output. For other groups, we recorded 48 h to obtain two technical runs per cage (day-1 and day-2), each summarized by 24 h AUC. Data were rewrapped to a standardized 24-hour format, aligning data from different cages to Zeitgeber time (ZT0 = lights on) to facilitate comparison of daily production patterns.

### *In vitro* profiling of strain metabolic activity

To assess individual metabolic capabilities, bacterial strains were grown from glycerol stocks in 3 mL of Mega medium or modified Chopped Meat medium for 72 hours at 37 °C under anaerobic conditions. Upon reaching stationary phase, cultures were harvested by centrifugation (5,000 × g for 10 min) and cell pellets were washed twice with an equal volume of pre-reduced, sterile phosphate-buffered saline (PBS) to remove residual growth media. The resulting cell pellets were resuspended in 0.75 mL of a modified SAAC minimal medium containing 20 amino acids. To assess specific metabolic pathways, phenol, 4-cresol, and 4-ethylphenol production was profiled by supplementing the SAAC medium with 20 µM of their respective precursors: 4-hydroxybenzoic acid, 4-hydroxyphenylpyruvic acid, and 4-vinylphenol. All other metabolites were profiled in non-supplemented SAAC media. Following a 96-hour incubation at 37 °C, cultures were stored at –80 °C. Metabolite concentrations, including aromatic amino acids and short-chain fatty acids (SCFAs), were subsequently quantified via LC-MS.

### Creatinine measurement

Creatinine concentrations were primarily measured using a colorimetric creatinine assay kit (Abcam #ab204537), following the manufacturer’s protocol. In cases where urine volume was limited, creatinine levels were instead quantified by LC-MS. LC-MS–based measurements showed strong correlation with values obtained from the colorimetric assay (data not shown).

### Metagenomic sequencing

Bacterial isolates and synthetic communities were processed using a unified sequencing workflow. Cell pellets were obtained by centrifugation under anaerobic conditions. Genomic DNA was isolated using the DNeasy PowerSoil HTP Kit (Qiagen), and DNA concentrations were quantified in 384-well plates using the Quant-iT PicoGreen dsDNA Assay Kit (Thermo Fisher). Library preparation was conducted in 384-well format using a miniaturized protocol adapted from the Nextera XT method (Illumina). DNA input was normalized to 0.18 ng/μL using a Mantis liquid handler (Formulatrix). Samples with lower concentrations were not diluted further. Tagmentation, neutralization, and PCR amplification were performed using the Mosquito HTS (TTP Labtech), with a final reaction volume of 4 μL per sample. Custom 12-bp dual unique indexing primers were used during PCR to prevent barcode misassignment associated with patterned flow cells. Libraries were pooled at specified molar ratios and purified with Ampure XP beads (Beckman Coulter) to remove contaminants and size-select fragments between 300 bp and 1.5 kb. Final library pools were assessed for quality and concentration using a Fragment Analyzer (Agilent) and quantitative PCR (Bio-Rad). Sequencing was carried out on either the NovaSeq S4 or NextSeq High Output platforms (Illumina), with 2×150 bp read lengths. We aimed to obtain 5–10 million paired-end reads per bacterial isolate and 20–30 million reads per synthetic community sample.

### Metagenomic mapping

Paired-end sequencing reads were aligned to the hCom2a reference genome database using Bowtie2 (v2.3.7) with the following parameters: --very-sensitive, --maxins 3000, -k 300, --no-mixed, --no-discordant, --end-to-end, and --no-unal. These settings enforced highly sensitive global alignments while suppressing unpaired, discordant, and unaligned reads. The alignment output in SAM format was streamed directly into Samtools (v1.9) for conversion to BAM format, followed by coordinate-based sorting and indexing.

Taking advantage of the fact that every organism in the hCom2 database has been carefully assembled, we used a highly sensitive pipeline, NinjaMap, to translate read alignment in the above BAM file into community composition, including detection of very low abundance organisms (<10^-6^). The complete metagenomic read analysis and the specific read mapping pipeline are available at https://github.com/FischbachLab/nf-ninjamap. To maximize accuracy and mitigate errors arising from imperfect database genomes in this study, we masked regions in NinjaMap based on mis mapped regions identified from the dropout strains in the dropout experiments. Reads that map to these masked regions in the subsequent analyses are assigned directly to the escrow bin.

To ensure high-confidence alignments, only reads with ≥99% identity across their full length were retained. On average, 4.95% of reads (range: 4.10%–8.35%) did not align to the reference. To investigate the source of these unaligned reads, a representative subset was queried against the NCBI nucleotide (nt) database using BLAST+ (v2.11.0) via the ncbi/blast:latest Docker image. The following parameters were applied: -outfmt ‘6 std qlen slen qcovs sscinames staxids’, -dbsize 1000000, and -num_alignments 100.

Top BLAST hits were defined as alignments with e-values ≤ 1e–30, percent identity ≥ 99%, and bit scores within 10% of the best hit for each query. Taxonomic summaries were generated using the ktImportTaxonomy script from the Krona toolkit, with default settings. Reads were aggregated by NCBI Taxonomy ID and genus. Most hits corresponded to taxa closely related to members of the synthetic community, while a portion mapped to the mouse genome. These results suggest a negligible level of environmental or cross-sample contamination in our sequencing data.

### Assessment of the ecological effects of strain dropouts

Variability in colonization complicates interpretation of ecological responses to strain removal (Fig. 1d). For some strains, large differences from the mean are indistinguishable from stochastic variation; for others, even small shifts are significant and potentially important. To account for this variability, we quantified each strain’s response relative to its typical behavior using a Z-score–based approach. For each strain in a dropout community, we first estimated its baseline abundance and variability from all other communities in the same batch (excluding the corresponding dropout itself) and then calculated a Z-score describing its deviation in the dropout community (**fig. s2a**). This normalization highlights context-specific responses while down-weighting fluctuations driven by unstable colonization. For example, a modest but consistent increase in a stable colonization (e.g., Strain A in Δ3, **fig. s2a**) can be more informative than a large fluctuation in a strain that colonizes inconsistently (e.g., Strain B in Δ2, **fig. s2a**).

### Assessment of genome-encoded acetate production capacity

Genome-encoded acetate-producing capacity was assessed by KEGG orthology (KO) assignment using KofamScan v1.3^34^ against KEGG KOfam HMM profiles^35^. A strain was classified as “acetate-from-EMP capable” if it encoded a complete or near-complete EMP module (≥75% of ten steps, with isoenzyme alternatives considered functionally equivalent), at least one pyruvate-dissipating enzyme (PFOR, PflB, or PDH), and both phosphotransacetylase (Pta) and acetate kinase (AckA). Wood-Ljungdahl pathway presence required acetyl-CoA synthase (AcsB; K14138) and formate-tetrahydrofolate ligase (Fhs; K01938), validated by tblastn against Ammonifex degensii *AcsB*. Per-strain calls are in **Table. S4**.

BLAST analysis was performed using TBLASTN (version 2.14.0+) to identify genomic sequences homologous to the genes of interest.

### Sample preparation for metabolomics

Mouse cecal contents and urine samples were processed for untargeted metabolomics analysis. For cecal samples, wet tissue (20–40 mg) was weighed into 2 mL screw-cap tubes preloaded with six 2.3 mm ceramic beads (RPI research products 9839). An extraction solvent consisting of 40:40:20 methanol: acetonitrile: water (v/v/v), containing 5 μM 4-chlorophenylalanine as an internal standard, was added at a volume of 20 μL per mg of tissue. Samples were homogenized using a QIAGEN TissueLyser II at 25 Hz for 10 minutes. Following homogenization, samples were centrifuged at 18,200 × g for 15 minutes at 4 °C. The resulting supernatants were collected, and 5 μL was injected for liquid chromatography–mass spectrometry (LC-MS) analysis. For mouse urine samples, 5 μL of urine was extracted with 100 μL of methanol containing 5 μM 4-chlorophenylalanine as an internal standard. Samples were incubated on ice for 20 minutes to facilitate protein precipitation, then centrifuged at 18,200 × g for 15 minutes at 4 °C. The supernatant was collected, and 5 μL was injected for LC-MS analysis.

Mouse gallbladder and small intestine samples were extracted for bile acid analysis by LC–MS. Gallbladder contents (2 µL) were extracted in 100 µL methanol containing 5 µM d^4^-cholic acid as an internal standard. Samples were vortexed briefly and centrifuged at 18,200 × g for 10 min at 4 °C. The supernatant was diluted 10-fold in methanol prior to transfer to LC–MS vials for analysis.

For small intestine measurements, in some experiments, only a portion of the small intestine was extracted for bile acid analysis in order to preserve remaining tissue for parallel metagenomic sequencing. In these cases, regional bile acid measurements were used to assess local bile acid concentrations (reported in µM) rather than total intestinal bile acid pool size (reported in µmol). To measure total bile acid pool, the entire small intestine (including duodenum, jejunum, and ileum) was transferred to a 5 mL Falcon tube and homogenized in 3 mL 40:40:20 methanol: acetonitrile: water (v/v/v) with 5 µM d^4^-cholic acid with 6.5 mm ceramic beads (Omni 19-682) using a QIAGEN TissueLyser II (25 Hz, 10 min). Homogenates were centrifuged at 18,213 × g for 15 min at 4 °C, and the resulting supernatants were diluted 100-fold in methanol before LC–MS analysis. For regional bile acid measurements, approximately 100 mg of small intestine tissue (jejunum) was weighed into a 2 mL microcentrifuge tube and homogenized in 40:40:20 methanol:acetonitrile:water (v/v/v) at a 100-fold dilution. After clarification by centrifugation, extracts were transferred directly to LC–MS vials for analysis.

SCFA Derivatization and Quantification: SCFAs were quantified following 3-nitrophenylhydrazine (3-NPH) derivatization as previously described^36^. Briefly, 20 μL of the cecal supernatant was reacted with 20 μL of a derivatization cocktail (12 mM EDC, 21 mM 3-NPH, and 2.4% v/v pyridine in methanol) at 4 °C for 1 hour. The reaction was quenched with 200 μL of 0.5 mM β−mercaptoethanol in water. After centrifugation (18,200 x g, 10 min), the final supernatant was transferred to vials for LC-MS analysis.

### Liquid chromatography/mass spectrometry

Bile acids and SCFAs were analyzed using an Agilent 6530 quadrupole time-of-flight (Q-TOF) mass spectrometer coupled to an Agilent 1290 Infinity II UPLC system. A dual Agilent Jet Stream electrospray ionization (AJS-ESI) source was used under extended dynamic range (m/z up to 1700). AJS-ESI source settings were as follows: gas temperature, 300 °C; drying gas flow, 7.0 L/min; nebulizer pressure, 40 psig; sheath gas temperature, 350 °C; sheath gas flow, 10.0 L/min; capillary voltage (VCap), 3500 V; nozzle voltage, 1400 V; and fragmentor voltage, 200 V.

Bile acids were separated by a Kinetex C18 column (1.7 μm, 2.1 × 100 mm; Phenomenex). Mobile phase A consisted of 0.05% formic acid in water, and mobile phase B consisted of 0.05% formic acid in acetone. 5 μL of each sample was injected via autosampler. Chromatographic separation was achieved using the following gradient at a flow rate of 0.35 mL/min: 25% B at 0–1 min; linear increase to 75% B at 25 min; ramp to 100% B at 26 min; held at 100% B until 30 min; and re-equilibrated to 25% B at 32 min.

SCFAs were separated using an Acquity UPLC BEH C18 column (2.1 mm × 100 mm, 1.7 μm particle size, 130 Å pore size; Waters, Milford, MA). The mobile phases were water (A) and methanol (B), with a flow rate of 200 μL/min. The LC gradient was as follows: 0–1 min, 10% B; 1–5 min, increase to 30% B; 5–11 min, ramp to 100% B; 11–14 min, hold at 100% B; 14.5–22 min, re-equilibrate to 10% B. The autosampler temperature was maintained at 5 °C, the column temperature at 60 °C, and the injection volume was 10 μL. The m/z values for derivatized SCFAs (in negative ion mode) were acetate: 194.0571; propionate: 208.0728; butyrate / Isobutyrate: 222.0884; valerate / isovalerate / 2-methylbutyrate: 236.1041. Aromatic amino acids derivatives were analyzed using a Thermo Q Exactive HF Hybrid Quadrupole-Orbitrap mass spectrometer coupled to a Thermo Vanquish UPLC system. MS1 scans were acquired over an m/z range of 60–900 at a resolution of 60,000. Data was collected in centroid mode with a loop count of 4 and an isolation window of 1.2 Da. Chromatographic separation was performed using an Acquity UPLC BEH C18 column (2.1 mm × 100 mm, 1.7 μm particle size, 130 Å pore size; Waters, Milford, MA). A 5 μL aliquot of each sample was injected via autosampler. The mobile phases consisted of water (A) and methanol (B), both containing 0.1% formic acid. Separation was achieved at a flow rate of 0.35 mL/min using the following gradient: 0 min, 0.5% B; 4 min, 70% B; 4.5–5.4 min, 98% B; 5.6 min, re-equilibrated to 0.5% B.

