## Supplementary information for "A single-strain dropout screen reveals mechanistic links between microbial ecology and metabolism"

### SUPPLEMENTARY TEXT

#### Strain dropouts influence community-level metabolism

*Propionate production*

Our attempt to drop out *Clostridium leptum* was one of the 12 that failed (Δ**Cl,* **fig. s3h**). However, in this community, we observed an unexpected result: Δ**Cl*-colonized mice had elevated levels of propionate (**fig. s3g**). Further analysis of the metagenomic sequencing data revealed an increase in the relative abundance of several strains including *Megasphaera, Faecalibacterium prausnitzii,* and *Ruminococcus bromii*. We cross-referenced BLAST annotations of propionate-producing enzymes, particularly enzyme *lcdA* in the acrylate pathway, and found that *Megasphaera* stood out as both a known propionate producer and one of the strains that increased in Δ**Cl* (Z = 2.10) (**fig. s3e-h**). We suspect *C. leptum* was present at a greatly reduced (but nonzero) level in the inoculum through contamination of one of the strain stocks. This change in the inoculum condition may have disrupted early colonization dynamics, allowing *Megasphaera* and others to expand. Consistent with this view, Δ*Fplau,* the community with the lowest propionate levels in our screen, is associated with significantly decreased *Megasphaera* relative abundance (Z = -2.09) (**fig. s3i-j**). These findings suggest that even a reduction in the relative abundance of a keystone strain like *C. leptum* can have a long-lasting impact on community structure and metabolic output.

*Butyrate production*

In one of our hCom2a colonized mice, we observed marked differences in SCFA profiles between two cages, despite using the same inoculum. In one cage, butyrate levels increased dramatically from ~1,000 μM to ~6,000 μM, while acetate levels decreased from ~13,000 μM to ~10,000 μM (**fig. s3l**). We hypothesized that a strain capable of converting acetate to butyrate may have colonized differentially between the two cages, leading to this shift in SCFA composition (**fig. s3k**). To explore this possibility, we performed a computational search for the butyryl-CoA:acetate CoA-transferase pathway, identifying 11 candidate strains harboring this function (**fig. s3e**). Among these, *F. prausnitzii* was one of the most significantly enriched strains in the high-butyrate cage (**fig. s3m**). These findings suggest that stochastic differences in colonization—particularly of a keystone metabolic strain like *F. prausnitzii*—can lead to profound shifts in community-level metabolic output, even under controlled inoculation conditions.

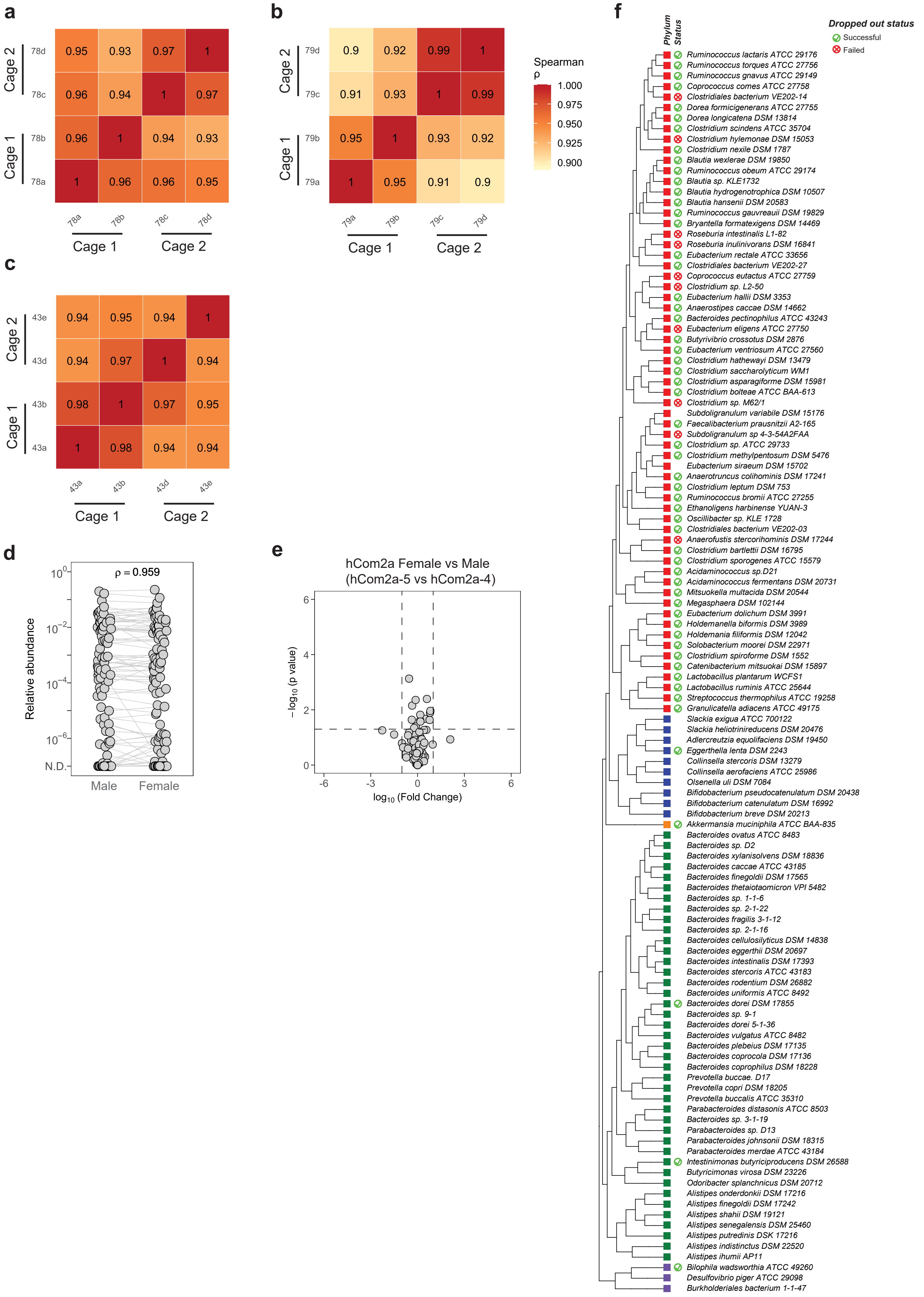
a
Cage 2
Cage 1
78d
78c
78b
78a
0.95 0.93 0.97 1
0.96 0.94 1 0.97
0.96 1 0.94 0.93
1 0.96 0.96 0.95
78a
78b
78c
78d
Cage 1
Cage 2
b
Cage 2
Cage 1
79d
79c
79b
79a
0.9 0.92 0.99 1
0.91 0.93 1 0.99
0.95 1 0.93 0.92
1 0.95 0.91 0.9
79a
79b
79c
79d
Cage 1
Cage 2
Spearman
ρ
1.000
0.975
0.950
0.925
0.900
c
Cage 2
Cage 1
43e
43d
43b
43a
0.94 0.95 0.94 1
0.94 0.97 1 0.94
0.98 1 0.97 0.95
1 0.98 0.94 0.94
43a
43b
43d
43e
Cage 1
Cage 2
d
ρ = 0.959
Relative abundance
10^0
10^−2
10^−4
10^−6
N.D.
Male
Female
e
hCom2a Female vs Male
(hCom2a-5 vs hCom2a-4)
− log10 (p value)
log10 (Fold Change)
6
4
2
0
-6
-3
0
3
6
f
Phylum
Status
Dropped out status
Successful
Failed
Ruminococcus lactaris ATCC 29176
Ruminococcus torques ATCC 27756
Ruminococcus gnavus ATCC 29149
Coprococcus comes ATCC 27758
Clostridiales bacterium VE202-14
Dorea formicigenerans ATCC 27755
Dorea longicatena DSM 13814
Clostridium scindens ATCC 35704
Clostridium hylemonae DSM 15053
Clostridium nexile DSM 1787
Blautia wexlerae DSM 19850
Ruminococcus obeum ATCC 29174
Blautia sp. KLE1732
Blautia hydrogenotrophica DSM 10507
Blautia hansenii DSM 20583
Ruminococcus gauvreauii DSM 19829
Bryantella formatexigens DSM 14469
Roseburia intestinalis L1-82
Roseburia inulinivorans DSM 16841
Eubacterium rectale ATCC 33656
Clostridiales bacterium VE202-27
Coprococcus eutactus ATCC 27759
Clostridium sp. L2-50
Eubacterium hallii DSM 3353
Anaerostipes caccae DSM 14662
Bacteroides pectinophilus ATCC 43243
Eubacterium eligens ATCC 27750
Butyrivibrio crossotus DSM 2876
Eubacterium ventriosum ATCC 27560
Clostridium hathewayi DSM 13479
Clostridium saccharolyticum WM1
Clostridium asparagiforme DSM 15981
Clostridium bolteae ATCC BAA-613
Clostridium sp. M62/1
Subdoligranulum variabile DSM 15176
Faecalibacterium prausnitzii A2-165
Subdoligranulum sp 4-3-54A2FAA
Clostridium sp. ATCC 29733
Clostridium methylpentosum DSM 5476
Eubacterium siraeum DSM 15702
Anaerotruncus colihominis DSM 17241
Clostridium leptum DSM 753
Ruminococcus bromii ATCC 27255
Ethanoligens harbinense YUAN-3
Oscillibacter sp. KLE 1728
Clostridiales bacterium VE202-03
Anaerofustis stercorihominis DSM 17244
Clostridium bartlettii DSM 16795
Clostridium sporogenes ATCC 15579
Acidaminococcus sp.D21
Acidaminococcus fermentans DSM 20731
Mitsuokella multacida DSM 20544
Megasphaera DSM 102144
Eubacterium dolichum DSM 3991
Holdemanella biformis DSM 3989
Holdemania filiformis DSM 12042
Solobacterium moorei DSM 22971
Clostridium spiroforme DSM 1552
Catenibacterium mitsuokai DSM 15897
Lactobacillus plantarum WCFS1
Lactobacillus ruminis ATCC 25644
Streptococcus thermophilus ATCC 19258
Granulicatella adiacens ATCC 49175
Slackia exigua ATCC 700122
Slackia heliotrinireducens DSM 20476
Adlercreutzia equolifaciens DSM 19450
Eggerthella lenta DSM 2243
Collinsella stercoris DSM 13279
Collinsella aerofaciens ATCC 25986
Olsenella uli DSM 7084
Bifidobacterium pseudocatenulatum DSM 20438
Bifidobacterium catenulatum DSM 16992
Bifidobacterium breve DSM 20213
Akkermansia muciniphila ATCC BAA-835
Bacteroides ovatus ATCC 8483
Bacteroides sp. D2
Bacteroides xylanisolvens DSM 18836
Bacteroides caccae ATCC 43185
Bacteroides finegoldii DSM 17565
Bacteroides thetaiotaomicron VPI 5482
Bacteroides sp. 1-1-6
Bacteroides sp. 2-1-22
Bacteroides fragilis 3-1-12
Bacteroides sp. 2-1-16
Bacteroides cellulosilyticus DSM 14838
Bacteroides eggerthii DSM 20697
Bacteroides intestinalis DSM 17393
Bacteroides stercoris ATCC 43183
Bacteroides rodentium DSM 26882
Bacteroides uniformis ATCC 8492
Bacteroides dorei DSM 17855
Bacteroides sp. 9-1
Bacteroides dorei 5-1-36
Bacteroides vulgatus ATCC 8482
Bacteroides plebeius DSM 17135
Bacteroides coprocola DSM 17136
Bacteroides coprophilus DSM 18228
Prevotella buccae. D17
Prevotella copri DSM 18205
Prevotella buccalis ATCC 35310
Parabacteroides distasonis ATCC 8503
Bacteroides sp. 3-1-19
Parabacteroides sp. D13
Parabacteroides johnsonii DSM 18315
Parabacteroides merdae ATCC 43184
Intestinimonas butyriciproducens DSM 26588
Butyricimonas virosa DSM 23226
Odoribacter splanchnicus DSM 20712
Alistipes onderdonkii DSM 17216
Alistipes finegoldii DSM 17242
Alistipes shahii DSM 19121
Alistipes senegalensis DSM 25460
Alistipes putredinis DSK 17216
Alistipes indistinctus DSM 22520
Alistipes ihumii AP11
Bilophila wadsworthia ATCC 49260
Desulfovibrio piger ATCC 29098
Burkholderiales bacterium 1-1-47

**Figure s1: Validation of the single-strain dropout screen. Related to Figure 1.** (**a-c**) Spearman correlation heatmaps showing community composition similarity among individual mice colonized with the same inoculum but housed in two different cages. Panels illustrate examples of minimal cage effects (**a, c**) and a cohort with a stronger cage effect (**b**). (**d**) Median relative abundances of hCom2a strains in cecal content from male and female germ-free Swiss-Webster mice. Each dot represents one strain; dots within a column represent median abundances from 3–4 co-housed mice (one cage). No significant differences in strain abundances were detected between sexes. (**e**) Volcano plot of strain relative abundances comparing female versus male mice. Strains below the detection limit were assigned a relative abundance of $10^{-8}$. Significance threshold: $p < 0.05$ (Student's t-test). (**f**) Phylogenetic tree of strains in hCom2a, indicating whether single-strain dropouts were attempted and whether they were successful (green, successfully removed; red, attempted but unsuccessful; others, not attempted).

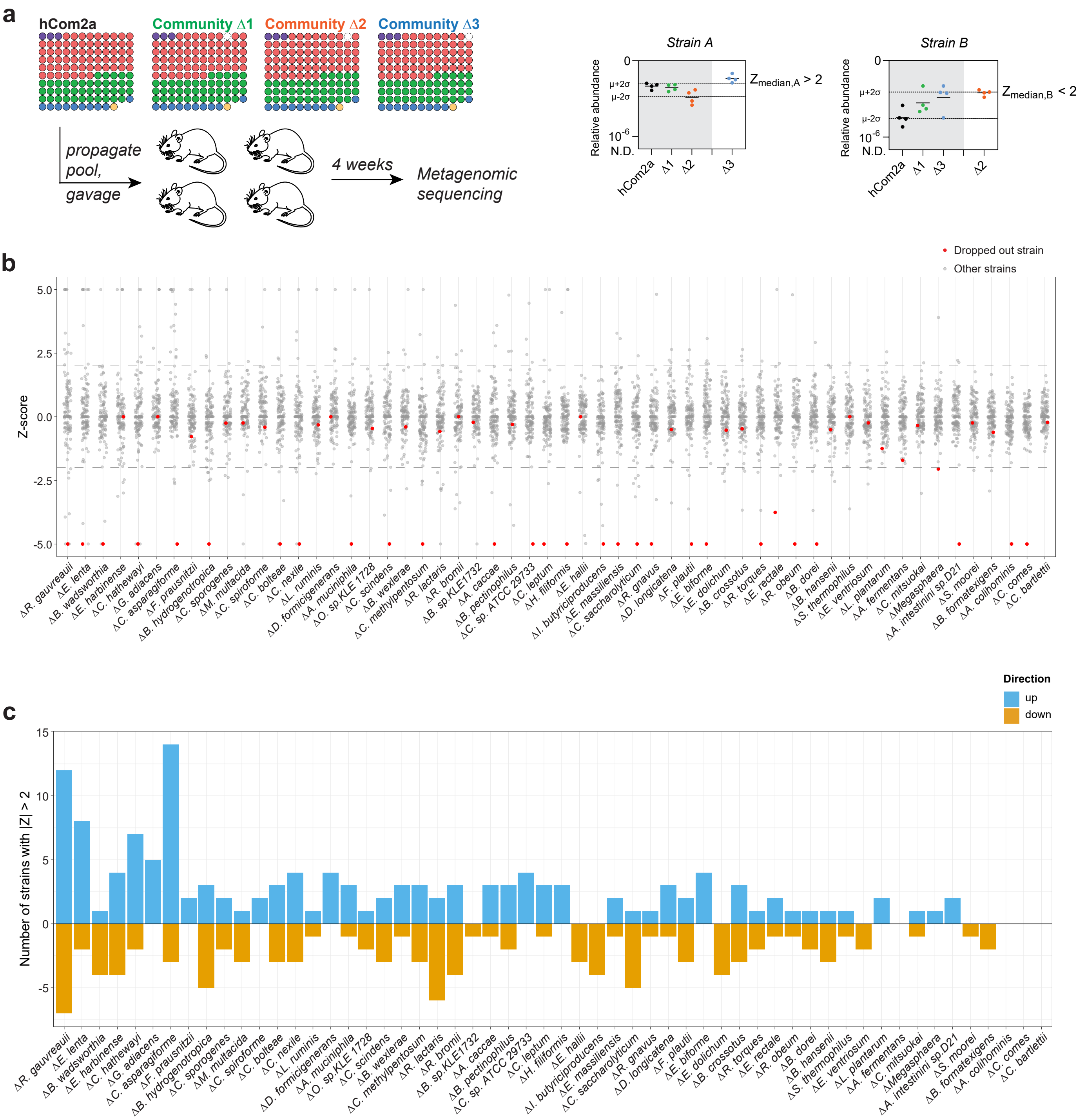
a
hCom2a
Community Δ1
Community Δ2
Community Δ3
propagate pool, gavage
4 weeks
Metagenomic sequencing
Strain A
Strain B
Relative abundance
0
μ+2σ
μ-2σ
$10^{-6}$
N.D.
hCom2a
Δ1
Δ2
Δ3
$Z_{median,A} > 2$
$Z_{median,B} < 2$
b
Dropped out strain
Other strains
Z-score
5.0
2.5
0.0
-2.5
-5.0
ΔR. gauvreauii
ΔE. lenta
ΔB. wadsworthia
ΔE. harbinense
ΔC. hathewayi
ΔG. adiacens
ΔC. asparagiforme
ΔF. prausnitzii
ΔB. hydrogenotropica
ΔC. sporogenes
ΔM. multacida
ΔC. spiroforme
ΔC. bolteae
ΔC. nexile
ΔL. ruminis
ΔD. formicigenerans
ΔA. muciniphila
ΔO. sp.KLE 1728
ΔC. scindens
ΔB. wexlerae
ΔC. methylpentosum
ΔR. lactaris
ΔR. bromii
ΔB. sp.KLE1732
ΔA. caccae
ΔB. pectinophilus
ΔC. sp. ATCC 29733
ΔC. leptum
ΔH. filiformis
ΔE. hallii
ΔI. butyriciproducens
ΔE. massiliensis
ΔC. saccharolyticum
ΔR. gnavus
ΔD. longicatena
ΔF. plautii
ΔE. biforme
ΔE. dolichum
ΔB. crossotus
ΔR. torques
ΔE. rectale
ΔR. obeum
ΔB. dorei
ΔB. hansenii
ΔS. thermophilus
ΔE. ventriosum
ΔL. plantarum
ΔA. fermentans
ΔC. mitsuokai
ΔMegasphaera
ΔA. intestinini sp.D21
ΔS. moorei
ΔB. formatexigens
ΔA. colihominis
ΔC. comes
ΔC. bartlettii
c
Direction
up
down
Number of strains with |Z| > 2
15
10
5
0
-5

**Figure s2: The ecological impact of single strain removal varies across communities. Related to Figure 2.** (**a**) A Z-score criterion was applied to assess whether changes in strain relative abundance were specifically associated with the removal of a strain. For each strain in each dropout community, the Z-score was calculated as follows: the mean and standard deviation of that strain's relative abundance were computed from all communities except the corresponding dropout, and these values were then used to calculate the Z-score for the dropout community. A small change in relative abundance can be meaningful if a strain colonizes consistently across mice (e.g., Strain A in Δ3), whereas a large change may be less significant when colonization is variable (e.g., Strain B in Δ2). (**b**) Ecological impact of strain removal. The effect of removing a strain on overall community architecture varied across dropout strains. The plot shows the Z-score of each strain in each of the 56 single-strain dropout communities. Red dot is the strain being dropped out in each strain dropouts. (**c**) Number of strains showing significant changes (Z-score > 2 or < –2) differed among dropout communities. The plot shows, for each of the 56 single-strain dropouts, the number of affected strains, excluding the intentionally removed strain from the count.

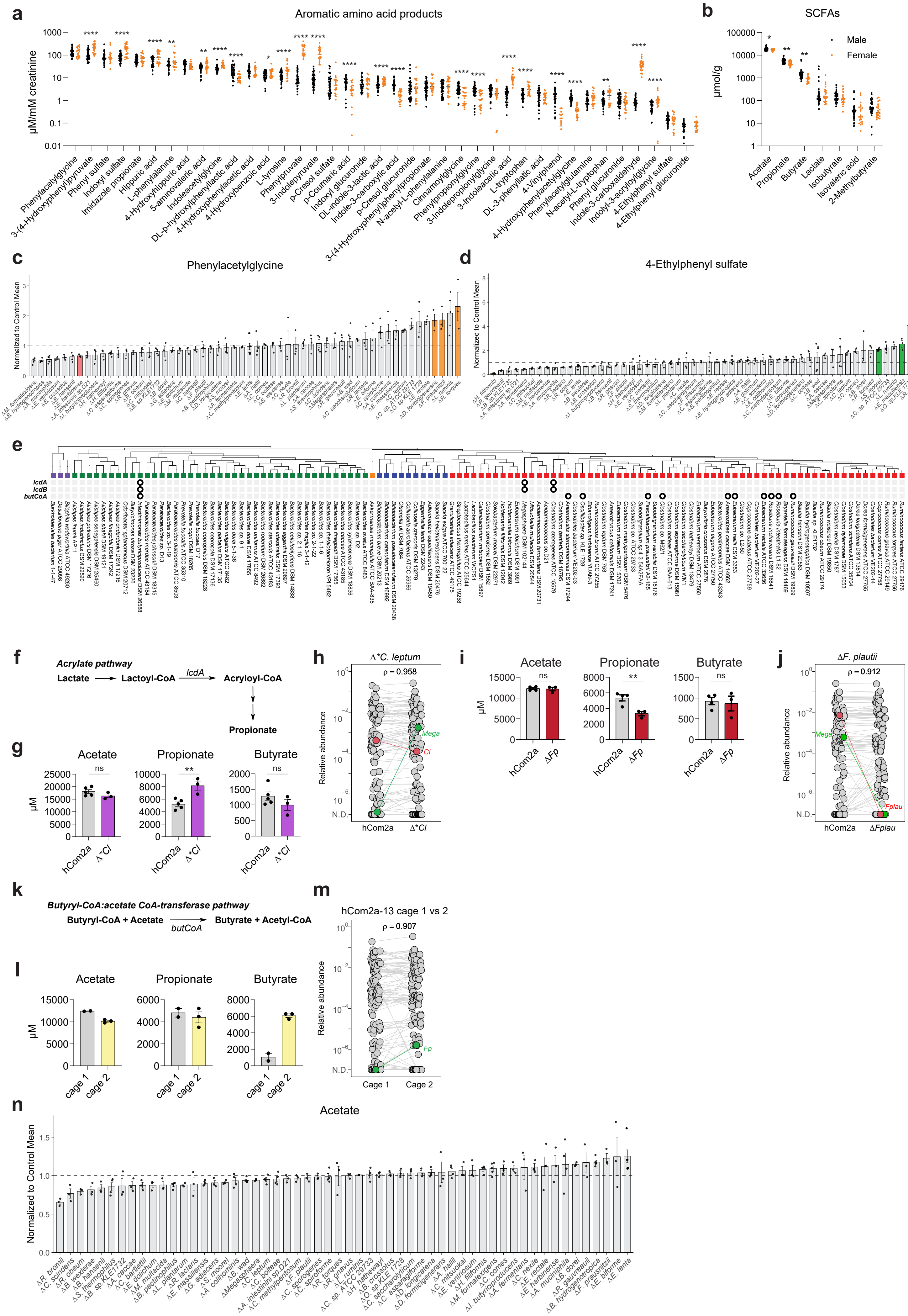

a
Aromatic amino acid products
μM/mM creatinine
Phenylacetylglycine
3-(4-Hydroxyphenyl)pyruvate
Phenyl sulfate
Indoxyl sulfate
Imidazole propionate
Hippuric acid
L-phenylalanine
4-Hydroxyhippuric acid
5-aminovaleric acid
Indoleacetylglycine
DL-p-hydroxylphenyllactic acid
4-Hydroxyphenylacetic acid
4-Hydroxybenzoic acid
L-tyrosine
Phenylpyruvate
3-Indolepyruvate
p-Cresol sulfate
p-Coumaric acid
Indoxyl glucuronide
DL-Indole-3-lactic acid
Indole-3-carboxylic acid
p-Cresol glucuronide
3-(4-Hydroxyphenyl)propionate
N-acetyl-L-phenylalanine
Cinnamoylglycine
Phenylpropionylglycine
3-Indolepropionylglycine
3-Indoleacetic acid
L-tryptophan
DL-3-phenyllactic acid
4-Vinylphenol
4-Hydroxyphenylacetylglycine
Phenylacetylglutamine
N-acetyl-L-tryptophan
Phenyl glucuronide
Indole-3-carboxaldehyde
Indolyl-3-acryloylglycine
4-Ethylphenyl sulfate
4-Ethylphenyl glucuronide
b
SCFAs
μmol/g
Male
Female
Acetate
Propionate
Butyrate
Lactate
Isobutyrate
Isovaleric acid
2-Methylbutyrate
c
Phenylacetylglycine
Normalized to Control Mean
d
4-Ethylphenyl sulfate
Normalized to Control Mean
e
lcdA
lcdB
butCoA
f
Acrylate pathway
Lactate
Lactoyl-CoA
lcdA
Acryloyl-CoA
Propionate
g
Acetate
Propionate
Butyrate
μM
hCom2a
Δ*Cl
h
Δ*C. leptum
ρ = 0.958
Relative abundance
Mega
Cl
N.D.
hCom2a
Δ*Cl
i
Acetate
Propionate
Butyrate
μM
hCom2a
ΔFp
j
ΔF. plautii
ρ = 0.912
Relative abundance
Mega
Fplau
N.D.
hCom2a
ΔFplau
k
Butyryl-CoA:acetate CoA-transferase pathway
Butyryl-CoA + Acetate
butCoA
Butyrate + Acetyl-CoA
l
Acetate
Propionate
Butyrate
μM
cage 1
cage 2
m
hCom2a-13 cage 1 vs 2
ρ = 0.907
Relative abundance
Fp
N.D.
Cage 1
Cage 2
n
Acetate
Normalized to Control Mean

**Figure s3: Strain removal impacts community-level metabolism. Related to Figure 3.** (**a**) Urinary levels of aromatic amino acid fermentation products in male and female germ-free Swiss-Webster mice colonized with hCom2a. Each dot represents an individual mouse. Significance was assessed using Student's *t*-test with false discovery rate (FDR) correction. $q < 0.05$(*), $q < 0.01$(**), $q < 0.001$(***), $q < 0.0001$(****). (**b**) Urinary levels of short-chain fatty acids (SCFAs) measured as in (**a**). (**c**) Waterfall plot of urinary phenylacetylglycine levels in each dropout community, normalized to the corresponding control group (N = 3–4 per condition). (**d**) Waterfall plot of urinary 4-ethylphenyl sulfate levels, as in (**c**). (**e**) Genomic distribution of several analyzed enzymes. Results of BLAST analysis for enzymes in production of propionate (*lcdA, lcdB*) and butyrate (*butCoA*). Black circles indicate strains with homologs detected. (**f**) Schematic of microbial propionate biosynthesis pathways. (**g**) Cecal SCFA levels in mice colonized with hCom2a versus $\Delta$**Cl* communities. (**h**) Relative abundance of individual strains in the $\Delta$**Cl* community. Each dot represents one strain; dots within a column represent median abundances from 3–4 co-housed mice (one cage). (**i**) Cecal SCFA levels in mice colonized with hCom2a versus $\Delta$*Fplau* communities. (**j**) Strain abundance analysis (as in (**h**)) for $\Delta$*Fplau.* (**k**) Schematic of the butyryl-CoA:acetate CoA-transferase pathway for butyrate synthesis. (**l**) Cecal SCFA levels in two cages colonized with the same hCom2 inoculum, illustrating cage-to-cage variation in SCFA profiles despite identical inoculation conditions. (**m**) Strain abundance analysis (as in (**h**)) for the two cages shown in (**j**). Expansion of a *Faecalibacterium prausnitzii* strain encoding a butyryl-CoA transferase gene may account for elevated butyrate production in cage 2. (**n**) A waterfall plot depicting the cecal acetate levels in each dropout community, normalized to its corresponding control group (N = 3–4 per condition). All graphs show mean +/- SEM. Statistical significance was determined using two-tailed Student's t-test except in (**a**) and (**b**); $p < 0.05$ (*), $p < 0.01$ (**), $p < 0.001$ (***).

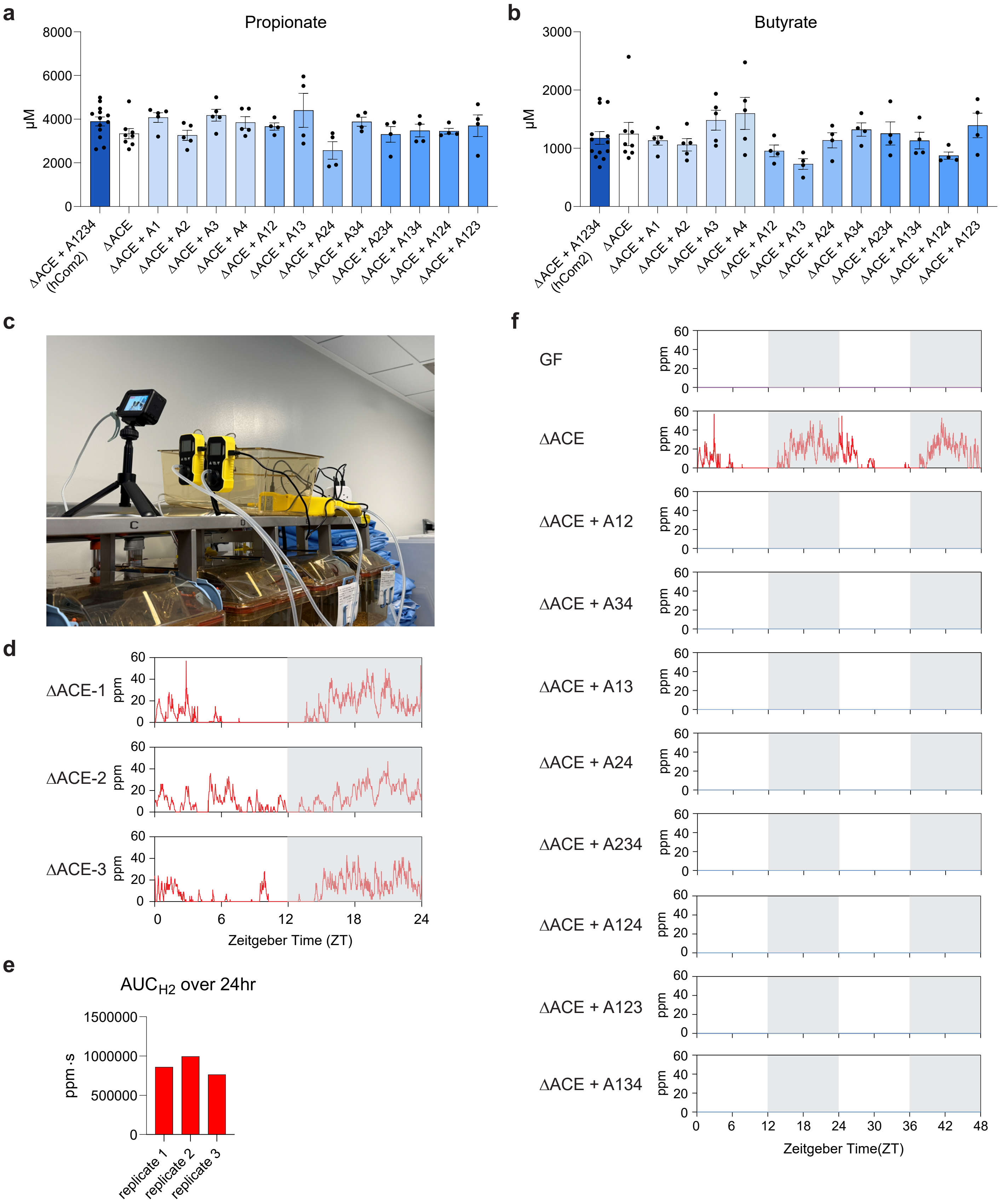
a
Propionate
µM
8000
6000
4000
2000
0
ΔACE + A1234 (hCom2)
ΔACE
ΔACE + A1
ΔACE + A2
ΔACE + A3
ΔACE + A4
ΔACE + A12
ΔACE + A13
ΔACE + A24
ΔACE + A34
ΔACE + A234
ΔACE + A134
ΔACE + A124
ΔACE + A123
b
Butyrate
µM
3000
2000
1000
0
ΔACE + A1234 (hCom2)
ΔACE
ΔACE + A1
ΔACE + A2
ΔACE + A3
ΔACE + A4
ΔACE + A12
ΔACE + A13
ΔACE + A24
ΔACE + A34
ΔACE + A234
ΔACE + A134
ΔACE + A124
ΔACE + A123
c
d
ΔACE-1
ΔACE-2
ΔACE-3
ppm
60
40
20
0
0
6
12
18
24
Zeitgeber Time (ZT)
e
AUC$_{H2}$ over 24hr
ppm·s
1500000
1000000
500000
0
replicate 1
replicate 2
replicate 3
f
GF
ΔACE
ΔACE + A12
ΔACE + A34
ΔACE + A13
ΔACE + A24
ΔACE + A234
ΔACE + A124
ΔACE + A123
ΔACE + A134
ppm
60
40
20
0
0
6
12
18
24
30
36
42
48
Zeitgeber Time(ZT)

**Figure s4: Systematic dissection of the acetogen niche in a defined community. Related to Figure 4.** (**a-b**) Targeted metabolic profiling of cecal propionate (**a**) and butyrate (**b**) concentrations in mice colonized with communities lacking all acetogens (ΔACE) or partially restored with specific acetogen groups. (**c**) Setup for continuous breath hydrogen gas measurement. Cage exhaust was routed to hydrogen sensors, with real-time data acquisition and visual monitoring performed over consecutive diurnal cycles. (**d**) Continuous 24-hour $H_2$ measurements from three biological replicates (communities gavaged to three independent groups of mice) of ΔACE. (**e**) AUC of $H_2$ production over 24 hr for the three biological replicates of ΔACE*.* (**f**) Continuous 48-hour $H_2$ measurements from cages colonized with germ-free (GF), ΔACE, and communities complemented with two or three groups of acetogens (ΔACE+A12 to ΔACE+A123). All graphs show mean +/- SEM.
